# Evolutionary Transitions in Heritability and Individuality

**DOI:** 10.1101/192443

**Authors:** Pierrick Bourrat

## Abstract

With a few exceptions, the literature on evolutionary transitions in individuality (ETIs) has mostly focused on the relationships between lower-level (particle-level) and higher-level (collective-level) selection, leaving aside the question of the relationship between particle-level and collective-level inheritance. Yet, without an account of this relationship, our hope to fully understand the evolutionary mechanisms underlying ETIs is impeded. To that effect, I present a highly idealized model to study the relationship between particle-level and collective-level heritability both when a collective-level trait is a linear function and when it is a non-linear function of a particle-level trait. I first show that when a collective trait is a linear function of a particle-level trait, collective-level heritability is a by-product of particle-level heritability. It is equal to particle-level heritability, whether the particles interact randomly or not to form collectives. Second, I show that one effect of population structure is the reduction in variance in offspring collective-level character for a given parental collective. I propose that this reduction in variance is one dimension of individuality. Third, I show that even in the simple case of a non-linear collective-level character, collective-level heritability is not only weak but also highly dependent on the frequency of the different types of particles in the global population. Finally, I show that population structure, because one of its effects is to reduce the variance in offspring collective-level character, allows not only for an increase in collective-level character but renders it less context dependent. This in turns permits a stable collective-level response to selection. The upshot is that population structure is a driver for ETIs. These results are particularly significant in that the relationship between population structure and collective-level heritability has, to my knowledge, not been previously explored in the context of ETIs.

## 1 Introduction

An evolutionary transition in individuality occurs when collective-level individuals emerge from the interactions of particle-level individuals during evolution. The topic of ETIs is intimately linked to one of the most important questions in evolutionary biology over the last 50 years, namely that of levels of selection (Williams, 1966; Sober and Wilson, 1998; Okasha, 2006; Godfrey-Smith, 2009; Wade, 2016). This link started to be particularly clear from the mid 90’s following the publication of Maynard-Smith and Szathmáry’s *The Major Transitions in Evolution* (Maynard Smith and Szathmary, 1995) and the the growing interest in ETIs (see Michod, 1999; Buss, 1983, 1987; Godfrey-Smith, 2009; Clarke, 2016a; Bouchard and Huneman, 2013; Calcott and Sterelny, 2011).^1^ For an ETI to occur, particle-level selection must in some sense shift to the collective-level. Yet, the notion of shift in selection is quite vague, and there is thus far, no fully satisfactory explanation for them. Yet, without a deep understanding of the nature of those ‘shifts’ in selection regimes from one level to the other, our hope to understand ETIs will be impeded. At the heart of ETIs thus lies the question of the levels of selection. But within the levels of selection question two problems are entangled. One concerns selection *sensu stricto*, that is considered independently from inheritance; the other concerns selection understood more broadly, that is including the questions relating to the response to selection and inheritance.

To see these two problems, it is useful to start from Lewontin’s famous three conditions for evolution by natural selection (Lewontin, 1970). Following Lewontin’s approach, an entity is a unit of selection, if it is part of a population that can evolve by natural selection (Godfrey-Smith, 2009). More specifically, the conditions say that evolution by natural selection will occur in a population in which there is 1) phenotypic variation, 2) this variation leads to differences in fitness, and 3) that this variation is heritable. As stressed by Lewontin, these three conditions can be satisfied at *any* level of organization, so that entities at different levels of organization can be ‘units of selection’. A natural move from there is to consider that to be a unit of selection is a necessary (but perhaps not sufficient) condition for Darwinian individuality^2^ so that the question of ETIs can be at least partly reduced to that of units of selection.^3^

We can see from Lewontin’s conditions that if selection *stricto sensu* – which can be defined as difference in fitness due to difference in phenotype – is important, so are inheritance and heritability – a population-level measure of inheritance. In fact, if the entities of a population exhibit differences in fitness without heritability, evolution might ensue, but natural selection will not be one of the evolutionary processes responsible for it. From this simple observation it is thus important to understand that the questions of how and in what sense both selection *stricto sensu* and heritability can shift from one level of organization to the other are equally important to solve the puzzle of the emergence of individuality in evolution. Yet, although the importance of the two questions has been expressed by several authors (e.g., Michod, 1999; Okasha, 2006; Herron et al, 2018; Griesemer, 2000), the question of the transitions in levels *of selection* has received much more attention than the question of the transitions in levels *of heritability*. For notable exceptions see Okasha (2006) and Herron et al (2018). The former question is undoubtedly an important one, and inasmuch as it still remains unresolved it deserves to be investigated. Yet, progress on the topic of ETIs will be impeded if no progress on transitions in levels of heritability is made.

In this paper, I aim at filling this gap. I provide an analysis of the evolution of heritability at different levels of organization in the context of ETIs. This analysis is different from both that of Okasha (2006) and Herron et al (2018). The former analyses heritability at different levels of organization from the perspective of the Price equation (as well as regression models derived from it) (Price, 1970, 1972) and uses Damuth and Heisler’s (1988) distinction between a conception of *collective* fitness, as the number of *particle* offspring produced, and a second conception of *collective* fitness as the number of offspring *collectives* produced. The former is often referred to as ‘multilevel selection 1’ and the latter as ‘multilevel selection 2.’ For reasons I cannot develop here, I believe the distinctions to be problematic in several respects (see Bourrat, 2015c, 2016, 2015b). I will thus depart from Okasha’s analysis.

Herron et al (2018) propose an analysis of heritability in the context of a type of ETIs in which collective offspring are genetic clones of a single parental collective and in which there is environmental variation. My model is different from theirs because I assume that offspring collectives can have multiple parents and consequently are not necessarily clones of their parent(s). Furthermore, although the effect of environmental variation is an important aspect of ETIs, I will not (or only briefly) consider it here. Rather my focus will be about the heritability of non-additive collective-level traits.

The results derived here apply to a maximally idealized system with the simplest and most mathematically tractable account of collective-level characters, and particle and collective generations. These results provide a baseline for a comparison. I will proceed in several steps. In Section 2, I present an additive model in which a collective character is the average character of its constituent particles, and in which offspring collectives have multiple parents. In this model collective-level heritability is strictly equal to particle-level heritability, whether or not there is a positive assortment between the offspring particles produced by a collective (a form of population structure). This leads me to consider, that in such a model, whether heritability is positive at the collective level is not, in and of itself, what permits to characterize the extent to which the collective level represents a ‘unit’ of evolution or a level of evolutionary individuality.

This leads me in Section 3 to argue that a low variance in average offspring-collective character produced by a parental collective tracks, at least partly, intuitions about whether the collective level exhibits individuality and is a more adequate criterion than the existence of collective-level heritability. I then show that a low variance in offspring-collective character from a parental-collective character can be achieved when the population structure increases. In Section 4, altering slightly the model presented in Section 2, I show that in a case of a non-linear collective trait, collective-level heritability is not only overall lower than particle-level heritability, but also highly context dependent when there is no population structure. Putting the different pieces together, I show in Section 5, that when population structure increases, collective-level heritability becomes less context dependent in the cases of non-linear collective traits considered.^4^ The upshot is that population structure is an important factor to consider in the context of ETIs, which is discussed in Section 6.

**Table 1:**
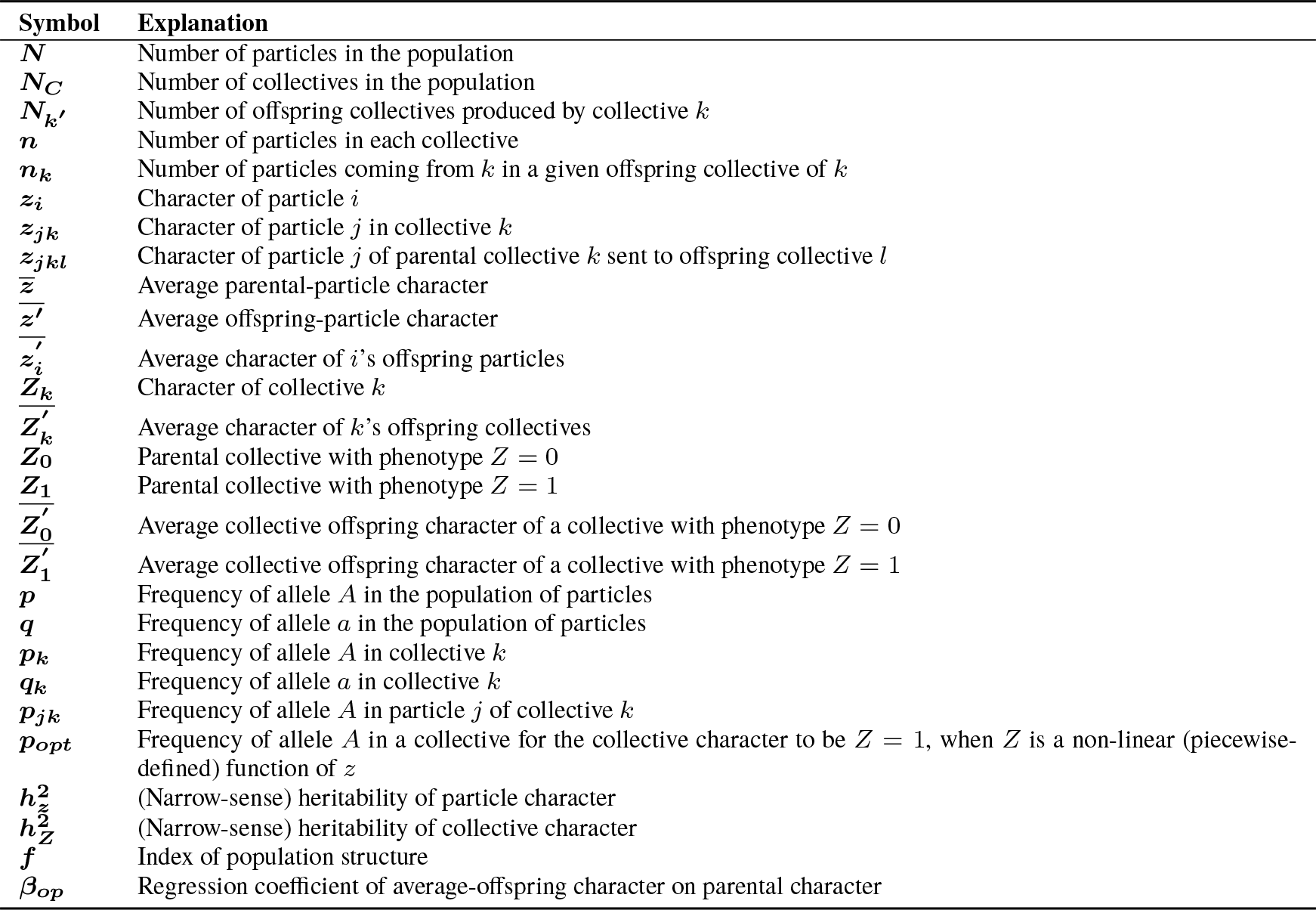
List of symbols

## 2 Collective-level Heritability and Additivity

Suppose a population of infinite size *N* made of haploid particles divided into an infinite number *N*_*C*_ of collectives, each of which is composed of *n* particles. The list of symbols used in the remainder of the article is reported in Table 1. I then assume that particles reproduce asexually, perfectly (more on this assumption in a moment) and simultaneously in discrete generations, and that particle and collective generations overlap perfectly – that is, collectives cease to exist when particles cease to exist and are reformed simultaneously with the offspring particles being produced. Note that because I am not directly interested in the difference made by selection in this article, I consider that all particles produce the same number of offspring particles at each generation. In other words, the particle character is neutral, so that each particle *i* has a character *z*_*i*_ which is independent from its fitness.

Let us now assume that a given collective *k* has a character *Z*_*k*_ which is equal to the mean particle character that composes it so that:

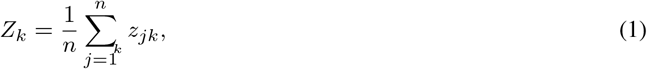

where *z*_*jk*_ is the character of particle *j* in collective *k*. We also assume that the character of each particle is genetically determined by one single locus with two alleles *A* and *a*, which have the respective frequencies *p* and *q* (with *p* + *q* = 1) in the global population of particles and *p*_*k*_ and *q*_*k*_ in the collective *k* (with *p*_*k*_ + *q*_*k*_ = 1). Since we assume that particles reproduce with perfect fidelity, we have:

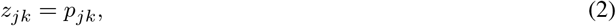

where *p*_*jk*_ is the frequency of allele *A* at the single locus of particle *j* in collective *k*. If the allele is *A*, we have *p*_*jk*_ = 1. If the allele is *a*, we have *p*_*jk*_ = 0. This leads to:

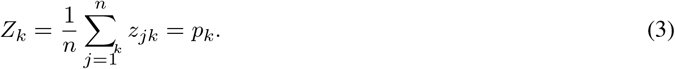

This model is effectively similar to the classical trait-group model first proposed by Wilson (1975), but with no difference in fitness and no interaction between the particles.

In this model, we can ask what the relationship between particle-level heritability 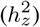 and collective-level heritability 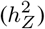 is. Before going further, I should make clear that I adopt a causal interpretation of heritability close to the one developed in Lynch and Bourrat (2017). Heritability represents, under my interpretation, the part of phenotypic variation that one can causally attribute to additive genetic variation – for that reason it refers to narrow-sense heritability (Falconer and Mackay, 1996). I consider that the relationship between two variables *X* and *Y* is causal if an ideal intervention on *X*, following Pearl (2009) and Woodward (2003), produces a difference in *Y*. In the case of heritability, if an intervention on the genetic composition of the population at the parental generation, produces a difference in the average offspring phenotype, then heritability is causal. Of course, many environmental variables are often correlated with the genetic composition of the population, the notion of ‘gene’ used when referring to heritability is not always the same in different context disciplines (e.g., molecular biology and evolutionary biology) (Lu and Bourrat, 2018), and there exist different ways to estimate heritability (see Falconer and Mackay, 1996, Chap. 10). All this means that heritability is fraught with a number of philosophical and theoretical problems, some of which have to do with the causal interpretation of heritability (see Godfrey-Smith, 2007, 2009; Downes, 2009; Bourrat, 2015a; Bourrat and Lu, 2017; Bourrat et al, 2017; Jacquard, 1983; Tal, 2009, 2012; Sesardic, 2005; Sarkar, 1998). I will consider here the simplest possible case in which there is no gene-environment correlation and interaction, so that the causal interpretation of heritability is unproblematic.

Narrow-sense heritability (*h*^2^) is defined as the ratio of *additive* genetic variance on total phenotypic variance (which can have additive genetic, non-additive genetic and environmental components). Starting with 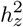, using the parent-offspring regression approach to heritability (see Falconer and Mackay, 1996, Chap. 10), computing this heritability is quite straightforward. In the additive case, with asexual organisms, particle-level heritability is expressed as:

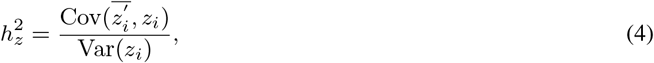

where 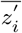 is the value of the average offspring character of entity *i*, 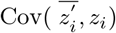 is the covariance between the average offspring character and the parental character, and Var(*z*_*i*_) is the variance of the parental character.

Since in our model, particles reproduce with perfect fidelity and there is no effect of the environment, we have:

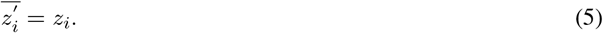

Therefore, recognizing that the covariance of a variable with itself is the variance of this variable, particle-level heritability can be rewritten as:

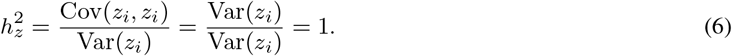

This result is not surprising: in the presence of variation in character, perfect inheritance of this character between parent and offspring without variation in the environment (or noise) is expected to be associated with maximal heritability.

The assumption of perfect fidelity in the context of measuring heritability might seem problematic to some. It would indeed be problematic if my aim was to characterize the effect of variation in the environment and noise at different levels of organization. As mentioned earlier, it is not. My aim instead is to study the effect of non-linear interactions between particles on collective-level heritability, as will become clear in Section 4.^5^ The environment and noise certainly have some important effects on heritability at different levels of organization as shown by Herron et al (2018), but adding these components of variance would greatly complexify the analysis provided here and would to some extent be orthogonal to my argument.

Moving on to collective-trait heritability 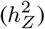, to be able to compute it, we first need to express the relationship between parental-collective character and average offspring-collective character in terms of particle distributions within parental and offspring collectives. To do so, I introduce the index *f*, which is an analog to Wright’s F-statistics (Wright, 1949; Weir and Cockerham, 1984). *f* is a measure of population structure from the point of view of particles. Note that *f* measures population structure assuming the size of collectives is given. This remark is important since population structure and collective size can be linked to one another in the following way: In the absence of population structure a population is a single large collective made of all the particles of the population since any particle has the same probability to interact with any other particle of the population. The existence of population structure makes the environment ‘viscous’ so that a given particle has a higher probability to interact with some particles than others, leading *de facto* to the formation of collectives or neighborhoods depending on the setting (Godfrey-Smith, 2008).

In contrast, when there is no population structure under my sense, that is *f* = 0, all particles form collectives (of a given fixed size) randomly. This means that an offspring particle of a parental collective has no more chance to form an offspring collective with another particle of the same collective than it has with any other offspring particle coming from other collectives. In a population of infinite size, *f* = 0 implies that one and only one particle coming from a given collective will be transmitted to a given offspring collective and consequently that each offspring collective has a number of parental collectives equal to its number of particles. If there is some population structure so that 0 < *f* < 1, an offspring particle has more chances to form an offspring collective with a particle coming from the same parental collective than it has with a particle coming from other collectives from the parental generation. In other words, on average, more than one particle is transmitted from a parental collective to each of its offspring collectives so that a given offspring collective can have more than one collective parent, but a lower number than when *f* = 0. Finally, when *f* = 1, offspring collectives are composed solely of the particle offspring of one parental collective. An illustration of the effect of population structure, as I define it, on the composition of offspring collectives is given in Figure 1.

**Fig. 1:**
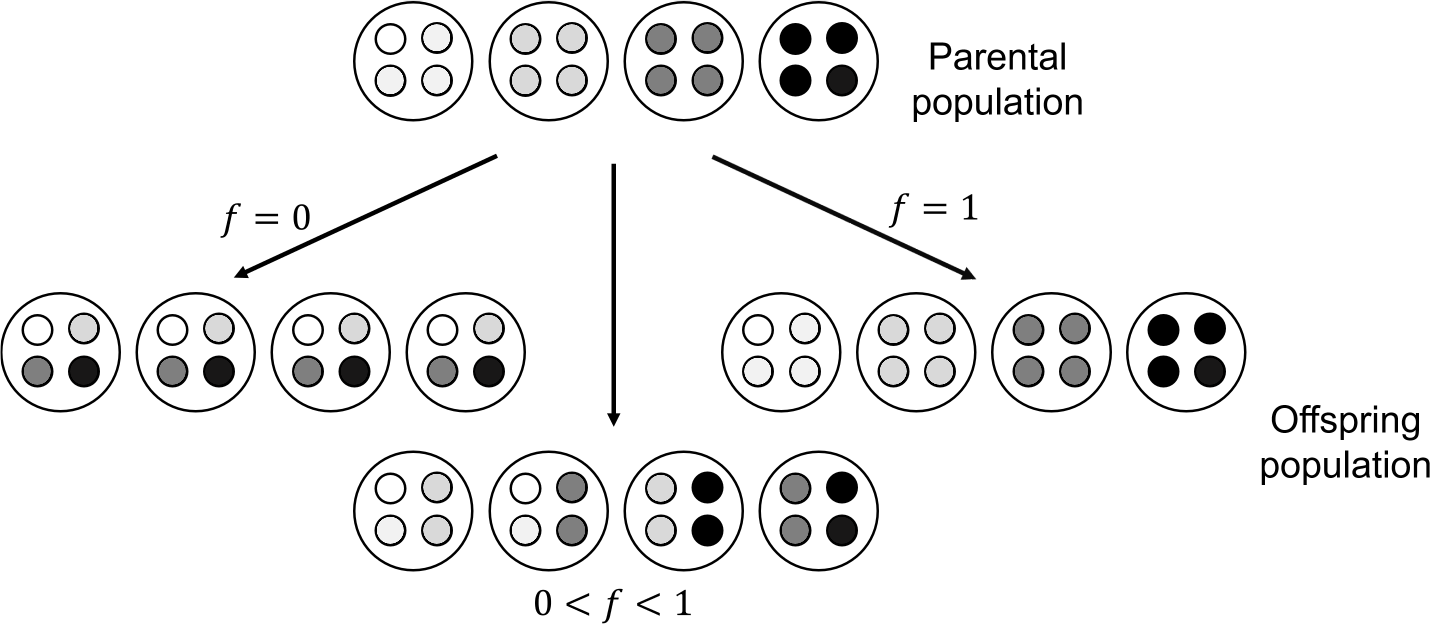
Illustration of the effect of *f* on the variance in offspring collective composition for a given parental collective. In each case, the parental origin of particles is tracked by their shade. Particles with the same shade in the offspring population originate from the same parental collective. When there is no population structure (*f* = 0, left) offspring particles coming from one parental collective have the same probability to be found in any offspring collective. As a result the composition of each offspring collective is the same. When there is some population structure (0 < *f* < 1, middle), offspring particles coming from one parental collective have more chances to be found in some offspring collectives rather than others. As a result, the composition of the four different collectives in the figure is different. When population structure is maximal (*f* = 1, right), offspring particles with the same parental collective have a probability of 1 to be found in the same offspring collective. Collectives reproduce perfectly.

From there, the value of average offspring-collective character from parental collective *k* can be calculated as the sum of two components modulated by the population structure *f*, namely one attributable to *k* in the absence of population structure (a single particle is sent to each of *n* offspring collectives produced by *k*) and another attributable to *k* when *f* is maximal (all particles *n* produced by *k* are sent to a single offspring collective). We have:

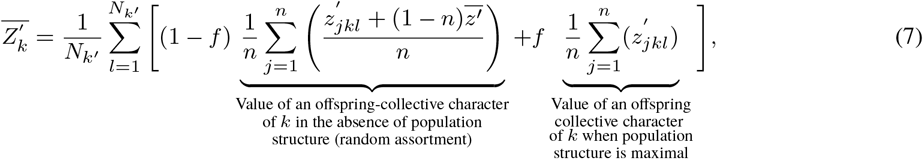

where *N*_*k*′_ is the number of offspring collectives produced by collective *k*, *l* is one offspring collective of collective *k*, 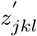 is the character of particle *j*’s offspring in collective *k*’s offspring *l*, and 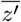 is the average offspring-particle character in the whole population. The index *f* can be rewritten as follows:

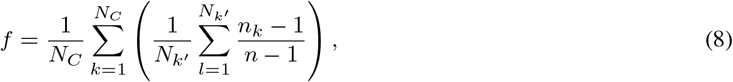

where *n*_*k*_ is the number of particles in offspring collective *l* coming from parental collective *k*. I will assume that parental collectives all contribute the same number of offspring particles to each of their offspring collectives so that we have:

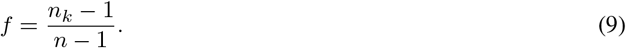

From equations (8) and (9), we can see that if only one particle offspring comes from a given parental collective in each offspring collective (which is what should be expected in a population of infinite size in which particles interact randomly) *f* is zero since *n*_*k*_ − 1 = 0. When all particles in a given offspring collective come from one given parental collective then *n*_*k*_ = *n*, in which case *f* = 1.

Since for any particle *i* we have *z*_*i*_ = *z*_*i*_, and because there is no difference in fitness between the different types of particles we have 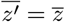, equation (7) can be rewritten as:

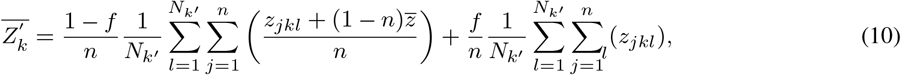

where *z*_*jkl*_ is the character of the particle *j* of parental collective *k* sent to offspring collective *l*. Developing (10) leads to:

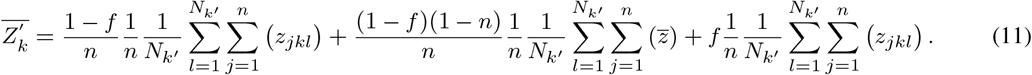

Recognizing that 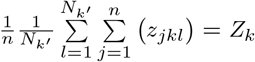 and 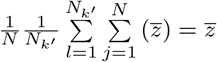, we can rearrange and simplify Equation (11) into:

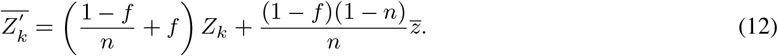

Since *Z*_*k*_ = *p*_*k*_ and 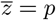, we can rewrite Equation (12) as:

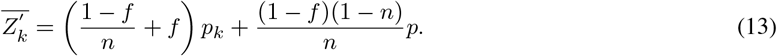

The first term of the right-hand side of Equation (13) 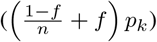 expresses the part of the collective character due to the structure of the population. The second term of the right-hand side 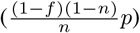 expresses the part of the collective character due to the random assortment of offspring particles in the formation of offspring collectives.

Generalizing for our model, the regression approach to heritability between one parent and the average offspring character in the case of sexual organisms (two parents) (Falconer and Mackay, 1996, Chap. 10), we have:

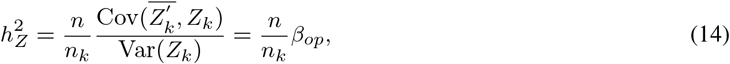

where 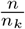 represents the number of parental collectives one given offspring collective has. When *f* = 0, that is when there is no population structure so that a parental collective sends only one particle per offspring collective, we have *n*_*k*_ = 1; when *f* = 1, that is when all the offspring particles of a collective are sent to a single offspring collective, we have 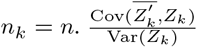 represents the slope of the best fitting line *β*_*op*_ when performing a one-parent-collective-average-offspring-collective regression.

If we now replace 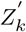 in (Equation (14) by its expression obtained in Equation (13), we get:

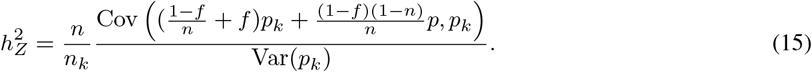

Using the distributive properties of variance, we can rewrite this equation as:

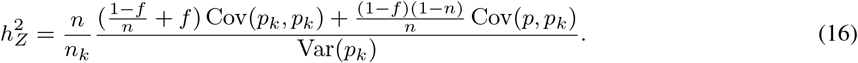

Since *p* is by assumption a constant and a covariance with a constant or between two constants is always nil, Equation (16) simplifies into:

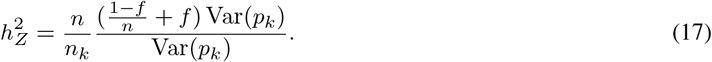

We can see that the second term of the right-hand side of Equation (17), namely 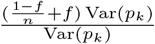 is simply 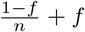 since 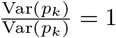. This leads to:

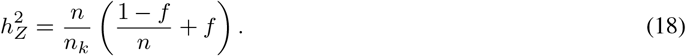

Recognizing that 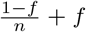 is always equal to 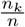, we thus have 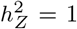 for any structure in the population and with collectives of any size.

The result obtained from Equation (18) shows us that in the case of a collective-additive trait, one can immediately derive its heritability at the collective level from the heritability of the particle trait from which it originates, whatever the population structure is, that is however the offspring particles of a parental collective are distributed in collective offspring. When the asexual particles reproduce perfectly, both 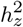 and 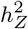 are equal to one. Although I do not show it here, when there is a normally distributed environmental deviation (or noise) of particle character centered around 0 that contributes additively to the collective character, the conclusion becomes that collective-level heritability is equal to particle-level heritability, even though it is inferior to one in both cases.

## 3 collective-level heritability and Collective Inheritance

In the previous section, I demonstrated that whatever level of assortment in the formation of offspring collectives between the offspring particles produced by a collective (measured by *f*), particle-level heritability and collective-level heritability of an additive-collective trait is always unity so long as the particles reproduce perfectly and there is no influence of the environment and no noise on the particle trait. Insomuch as collective-level heritability in one of the simplest possible models is derived directly from heritability at the lower level, surely the presence of collective-level heritability cannot be the sole criterion to consider when it comes to evaluate whether an entity is a unit of evolution or an individual in its own right with respect to inheritance.^6^ In fact, using this criterion has no discriminatory power since it would answer that any collective with an additive-collective trait is a unit of evolution.

In this Section, considering the limitations with collective-level heritability for characterizing a unit of evolution (or individuality), I put forward a different criterion with respect to inheritance for an entity to be a unit of evolution. I propose that the degree to which an entity is a unit of evolution is inversely proportional to the variance in offspring-collective-level characters it produces. In other words individuality, I suggest, amounts, at least partially, to the fidelity of character transmission from parental to offspring collective. Everything else being equal, a collective with a given collective-level character producing offspring with lower variance in collective-level character, scores higher on individuality than a collective producing offspring with a higher variance in collective-level character. One justification for this criterion comes from Maynard Smith and Szathmary (1995, p. 13) for whom an important feature of ETIs is the creation of new ways of transmitting information over time. Arguably, transmitting information can be regarded as equivalent to transmitting an entity’s character (at any level of organization) from one generation to the other with a high fidelity (which requires a low variance in offspring character, assuming the mean is close to that of the parental character).

As I show below, variation in the value of *f* can make important differences in the variance in offspring-collective character produced by a given collective. As such, if one accepts a low variance in offspring character produced by an entity as a criterion for individuality, increase in population structure represents an “engine” for ETIs, as it permits to increase collective-level heritability for non-additive characters. In Section 6, I will briefly mention what factors can contribute to increasing population structure during ETIs.

Let us start from the model presented in the previous section. Recall from Equation (3) that, by assumption, the collective character is proportional to the number of particles with allele *A* in the collective. For a given parental collective the variance in its offspring-collective character can be conceptualized as the outcome of two variances: First the variance originating from what this parental collective transmits to its offspring; Second, the variance transmitted to the offspring from the rest of the particles produced by other parents. We assume here that the parental contributions to a given offspring collective are independent. If we start from a case in which there is no population structure, a given parental collective transmits one and only one particle to each of its offspring collectives (recall that this is because the population has an infinite size); *n* − 1 offspring particles of its offspring collectives come from other parental collectives The allele of the transmitted particle depends on the composition of the parental collective. Its variance is that of a binomial law for a single trial and a probability *p*_*k*_ of transmitting *A* (*B* ~ (1, *p*_*k*_)). This variance is equal to *p*_*k*_*q*_*k*_. It is maximal when *p*_*k*_ is 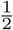. The distribution of the *n* − 1 particles coming from other collectives at the parental generation follows a binomial law for *n* − 1 trials and a probability of transmitting *A* equal to *p* (*B* ~ (*n* − 1, *p*)). The variance for this distribution is equal to (*n* − 1)*pq*.

Thus, when there is no population structure, under the assumptions made, we have a variance in offspring-collective character 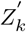 for a given parental collective with character *Z*_*k*_ equal to:

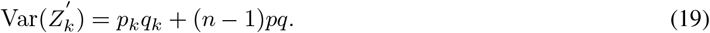

We can see two things from Equation (19). First, the value of this variance depends on the value of *p*_*k*_ and *p*. Second, unless *p*_*k*_ and *p* are both very large or very small, when there is no structure in the population, variance in collective-offspring character for a given parental collective will be high, and increases with *n*.

Suppose now that there is some structure in the population so that collectives transmit more than one offspring particle to their offspring collective. In such a case, a collective parent transmits *n*_*k*_ particles to its offspring and *n* − *n*_*k*_ particles of each offspring collective come from other collectives at the parental generation. The variance in the focal parental collective contribution follows a hypergeometric distribution of *n*_*k*_ draws in a collective of *n* particles with a number *p*_*k*_ of allele *A*.^7^ This variance is equal to 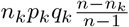. The variance in other parental collective contributions follows a binomial distribution of *n* − *n*_*k*_ trials and a probability *p* to transmit the allele *A* at each trial. We thus have:

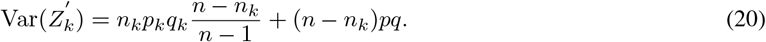

Assuming the extreme case where all the particles of a collective come from a single parent, we have *n*_*k*_ = *n*. Applying Equation (20), we get 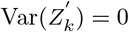. Note that when *n*_*k*_ = 1, Equation (20) becomes Equation (19).

We can now express Equation (20) in terms of *f*. From Equation (8) we can deduct that *n*_*k*_ is equal to:

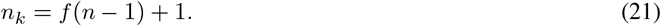

Replacing Equation (21) in Equation (20), we get:

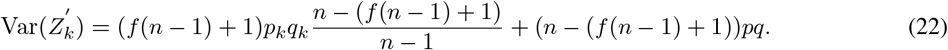

Equation (22) shows that the variance in collective offspring character for a parental collective depends both on the population structure (measured by *f*) and the size of collectives. The higher the collective size, the higher the variance in offspring-collective character, keeping *f* constant. The higher *f*, the lower the variance in offspring collective, keeping *n* constant. Furthermore, as *f* increases, the less 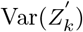 depends on the value of *p*_*k*_ and *p*. Finally, everything else being equal, when *p*_*k*_ < 0.5, the lower the value of *p*_*k*_, the lower the variance in collective offspring trait, and when *p*_*k*_ > 0.5, the higher the value of *p*_*k*_, the lower the variance in collective offspring trait.

If one dimension of individuality is the ability for an entity to reliably transmit the value of its character without too much variation, as I have suggested it is, then following my model, ETIs must have required some or the combination of three things. Namely, they must have required a population structure favoring a low number of particles in a collective, a much stronger assortment between particles coming from a parental collective than from any other collectives, and/or a low variance in parental collectives. These three factors —or a combination of them—will lead parental collectives to produce offspring collectives with the same character value as their parent.

It is interesting to note that in any real situation, everything else being equal, because the number of particles produced by a given collective is finite for a given parental collective, the higher the collective character variance in its offspring, the higher the number of offspring produced. This of course excludes cases in which the parental collective is genetically homogeneous and/or *f* = 1. This also assumes that the number of offspring particles transmitted by a parent is kept fixed. This gives scope for a trade-off between size and number of offspring collectives (which is modulated by *f*) when the collective trait is not neutral. I do not explore the consequences of this here, but leave it for future work.^8^

## 4 Heritability of Non-additive Collective Traits

I showed in Section 2 that when a collective trait is additive and particles reproduce perfectly, collective-level heritability is always equal to particle-level heritability 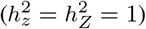. Given this result, it implies that collective-level heritability cannot be used to decide whether a collective is a unit since collective-level heritability is purely redundant with particle-level heritability. For that reason, I proposed instead in Section 3 that a better criterion for measuring the extent to which an entity is a unit or an individual is the ability for this entity (here the collective) to produce offspring which do not vary too much from their parent character. I further established the relationship between *f* and variance in collective offspring. Yet, so far I have only treated collective-level heritability for cases of additive collective-level character. I demonstrated that for such traits, *f* has no impact on collective-level heritability. In this and the next section, I extend my analysis for heritability of non-linear collective traits, that is collective-level traits that depend on the traits of the particles constituting a collective but that do not follow a linear function. I show that the conclusion reached in Section 2 cannot be extended to non-linear traits.

In this section, I demonstrate that collective-level heritability, at least for one sort of non-linear traits, is lower and more context-dependent than that of particle-level traits when offspring particles interact randomly to form offspring collectives. This result is significant insofar as it shows that heritability for non-linear collective traits is not redundant with particle-level heritability. One might thus consider that collective-level heritability can serve as a criterion for demarcating a collective-level unit or individual in the context of non-linear collective traits. A unit or individual following this criterion would be an entity that is part of a population in which there is a high level of collective-level heritability. Yet, as I show, collective-level heritability for non-linear collective traits is highly sensitive to allele frequencies. This means concretely that changing the allele frequencies in a population would force us to change on mind on the extent to which a collective is a individual. Rather, whether an entity is a individual, I claim, should be independent of the allele frequencies in the population. For that reason, the presence of collective-level heritability cannot, in and of itself serve as a reliable criterion for defining a unit of evolution or an individual. However, as I will show in Section 5, one way by which collective-level heritability can increase and be less context dependent is by increasing population structure (*f* > 0), which I proposed in Section 3 can be seen as a better criterion for individuality than the presence of collective-level heritability.

Non-linearity can be approached in different ways. A classical way is to consider that the character of a collective depends on a polynomial function of the characters of particles that compose the collective. Here I use a different notion, namely one in which the collective character is a piecewise-defined or hybrid function of particle character.

With a piecewise-defined function, the function’s domain is separated into different intervals over which a different (sub)function applies (Holtfrerich and Haughn, 2006, chap. 1). To see what I mean by that, take again the model presented in Section 2. This time, suppose that the character *Z* of a collective depends non-linearly on the proportion of particles with the allele *A* in the following way. We will assume here that *Z* is 1 when the proportion of particles with allele *A* within a collective has a certain frequency *P*_*opt*_, and 0 when this proportion is different from *P*_*opt*_. *Z* is thus defined as:

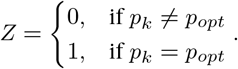

**Table 2:**
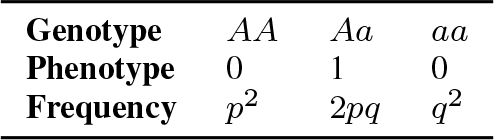
Genotype, phenotype and frequencies of two-particle collective genotypes at the Hardy-Weinberg equilibrium

**Table 3:**
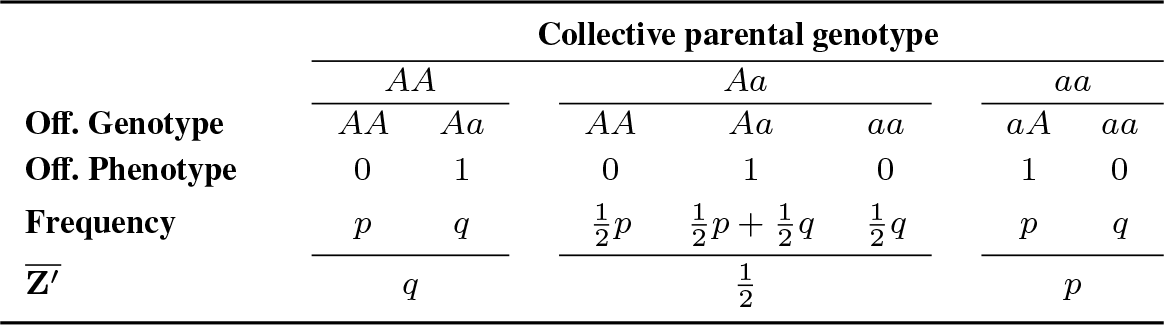
Frequencies of two-particle average offspring-collective phenotype for each parental-collective genotype at the Hardy-Weinberg equilibrium

In biological terms, this type of interaction could easily occur in *egalitarian* ETIs, during which two or more partners can now perform a function that none of them could perform independently (Queller, 1997), such as, for instance the synthesis of a protein.

collective-level heritability, in such cases, will depend on the different values of the parameters of the populations. To keep things simple, I present the cases in which collectives are made of two and four particles. I do not present the case for three-particle collectives because it is more complex than both the two- and four-particle cases. In fact, in the four-particle case some values of *f* which I will use can lead an equal number of particles to be systematically transmitted from one collective parent to all of its offspring collectives. In the case of three-particle collectives this is not possible. For example, although 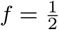 for the three-particle case means that, on average, a parental collective transmits two particles to its offspring, this necessarily implies, under my assumptions, that the collective sends half of the time one particle and half of the time all three particles to a given offspring collective. When there is variation in the number of offspring collectives produced by a parental collective, estimating the collective-level heritability from regressions becomes more complex. In the case of four-particle collectives, when 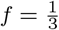, which is the example I will use in the next section, under my assumptions, collectives always send two particles to their offspring, so that the variance in contribution to offspring collectives is nil. A collective is, in such cases, an equal parent to all of its offspring, which makes the estimation of collective-level heritability easier. I start with the case of two-particle collectives.

When collectives are made of two particles, there exist three possible types of collectives, of which the frequencies follow the Hardy-Weinberg equilibrium (see Table 2) when the formation occurs from the random interaction of particles. Suppose now that only ‘heterozygote’ collectives (*Aa*) have a phenotype *Z* = 1 while the two ‘homozygote’ collectives have a phenotype *Z* = 0 (*AA* and *aa*). To calculate the heritability of the collective character, we first need to know the average offspring-collective character for the two possible parental collective phenotypes namely *Z* = 0 (*Z*_0_) and *Z* = 1 (*Z*_1_). This requires first computing the average offspring-collective phenotype of the three collective genotypes *AA*, *Aa*, and *aa*. These are reported in Table 3.

We then need to calculate the weighted average offspring-collective character of parental collectives with *Z* = 0 and *Z* = 1, which I symbolize by 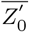 and 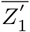 respectively. The value of 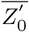 depends on the parental genotype frequencies of collectives *AA* and *aa*. From Table 2 and Table 3 we can compute the average offspring-collective character for these genotypes. It is given by the following equation:

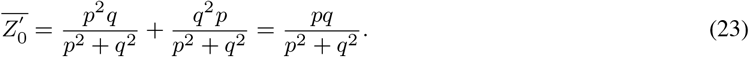

The value of 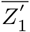 is found directly in Table 3 and is equal to:

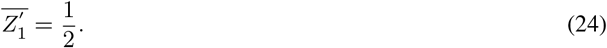

**Fig. 2:**
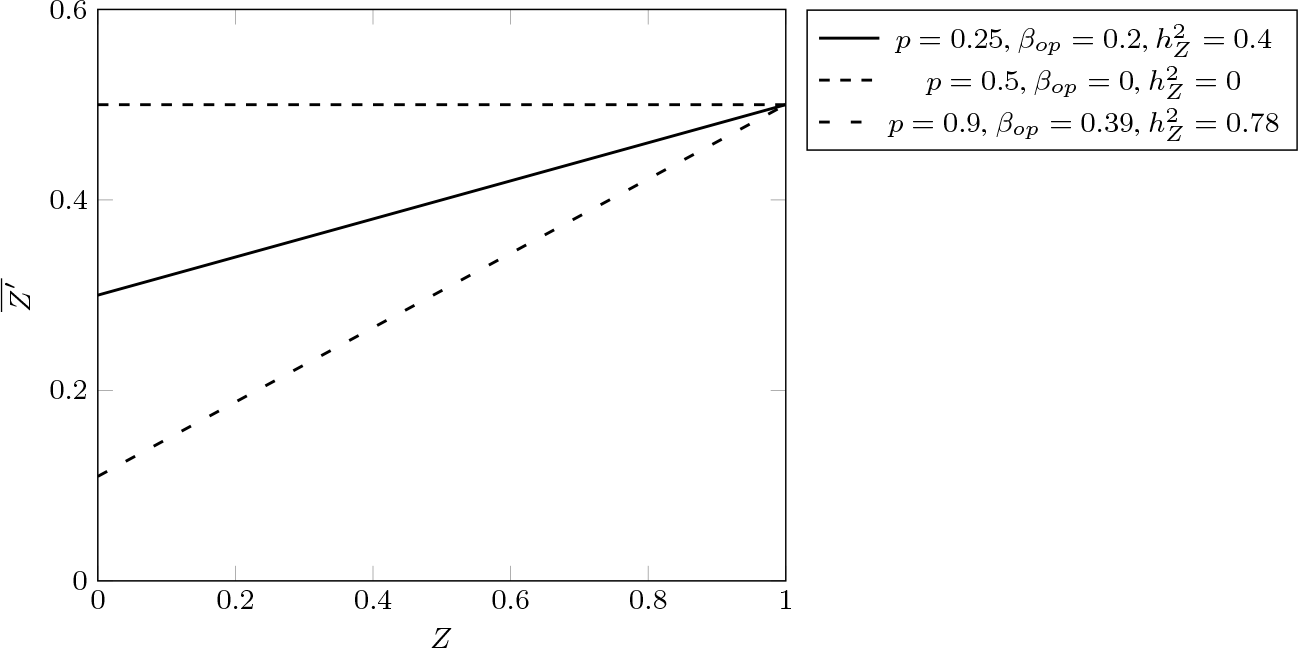
Linear regressions of average offspring-collective character on parental-collective character for the model of collectives with two-particles producing a non-linear trait in the absence of population structure (*f* = 0). Collective-level heritability is highly variable depending on the frequency *p* of allele *A*. *h*^2^ = 0.4 when *p* = 0.25, 0 when *p* = 0.5, and 0.78 when *p* = 0.9.

**Table 4:**
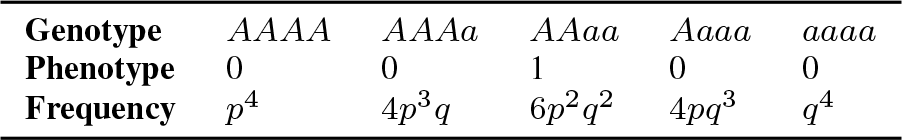
Genotype, phenotype and frequencies of four-particle parental collective-genotypes at the generalised Hardy-Weinberg equilibrium

With these two results, we can now plot the the average offspring character on parental character and find the slope of the best fitting line using the standard least-square method. These are shown in Figure 2 for three values of *p*, namely 0.25, 0.5, and 0.9.

As can be seen on Figure 2, when 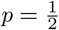, we have 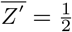 for parental collectives with a collective character *Z* = 0, and we also have 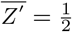 for parental collectives with a collective character *Z* = 1. The slope of the regression line of average offspring character is thus 0 and consequently collective-level heritability, calculated from Equation (14), is nil. Another observation is that the more distant *p* is from 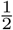, the larger collective-level heritability is. That said, it is always inferior to particle-level heritability 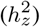. This observation can easily be explained. In a population in which most collectives are *AA*, these collectives become parents mostly of *AA* collectives due to the lack of variation in the population (*q* ≪ *p*), while *Aa* collectives become parents of collectives that are half of the time identical to them and almost half of the time *AA* or *aa* (*Z* = 0). Finally, *aa* collectives, with the same phenotype as *AA*, produce almost systematically offspring that are different from them, that is *Aa*. That said, they are so rare that they almost do not count in the weighted average offspring character of collectives with phenotype *Z* = 1. Thus, when *p* is very low or very high, the slope of the regression line tends toward 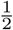. Since there are two collective parents per offspring collective, collective-level heritability (which is twice the slope of the regression line in this case) tends towards 1 but never reaches it.

Moving on to the four-particle setting, there are, in this case, five different possible collective genotypes with frequencies following the generalized Hardy-Weinberg equilibrium for two alleles. These are reported in Table 4. The average offspring-collective phenotype of the five possible collective genotypes *AAAA*, *AAAa*, *AAaa*, *Aaaa*, and *aaaa* are reported in Table 5. I suppose in this example that the genotype *AAaa* leads to the collective phenotype 1 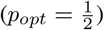 while all the other genotypes lead to the phenotype 0.

Using the same method as with the two-particle case presented earlier, we can calculate the weighted average offspring-collective character of parental collectives *Z*_0_. In this case it depends on parental genotype frequency of collectives *AAAA*, *AAAa*, *AAaa* and *aaaa*. The average offspring-collective character for these collectives can be calculated from Table 4 and Table 5. 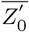 is given by the following equation:

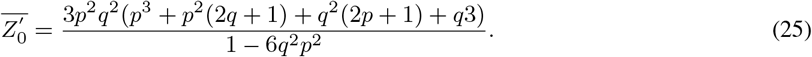

As with the two-particle-collective case presented earlier 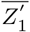, is found directly in Table 5 and is equal to:

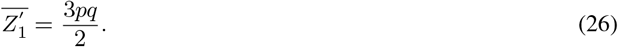

**Table 5:**
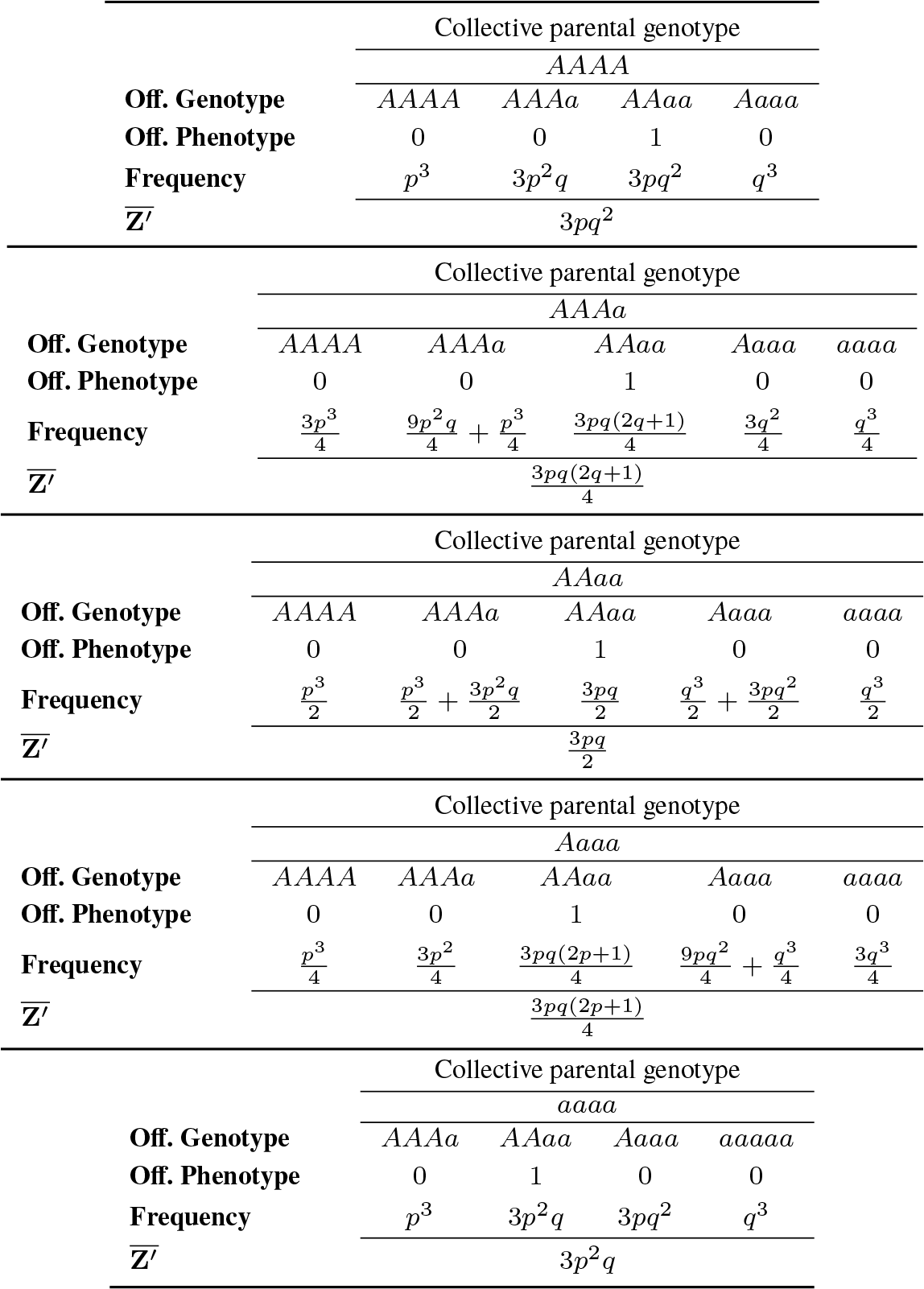
Frequencies of four-particle average offspring-collective phenotype for each parental-collective genotype with *f* = 0

**Fig. 3:**
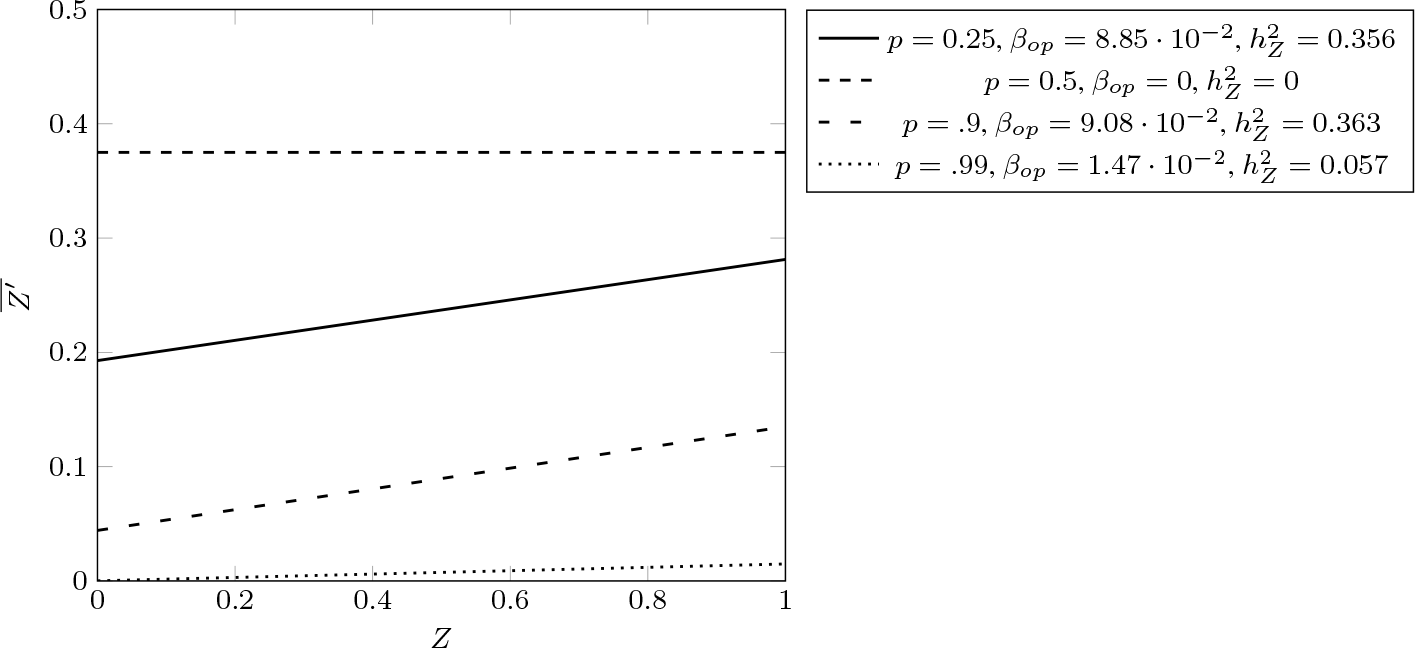
Linear regressions of average offspring-collective character on parental-collective character for the model of collectives with four particles producing a non-linear trait in the absence of population structure (*f* = 0). Collective-level heritability, like with the two-particle model (Figure 2) is highly variable depending on the frequency *p* of allele *A*. *h*^2^ = 0.356 when *p* = 0.25, 0 when *p* = 0.5, 0.363 when *p* = 0.9, and 0.057 when *p* = 0.99.

With these two equations, following the same method as previously, we can now plot the average offspring character on parental character and find the slope of the best fitting line *β*_*op*_ using the least-square method. These are shown in Figure 3 for four values of *p*: 0.25, 0.5, 0.9, and 0.99.

Following Equation (14), collective-level heritability is computed as four times the regression coefficient (*β*_*op*_) of average offspring-collective character on parental-collective character. The trend observed with the four-particle case is similar to that of the two-particle case. From Figure 3 we can see that 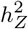 varies widely depending on the frequencies of the two alleles. Collective-level heritability never reaches the same value as particle-trait heritability (which, under my assumptions is always 1). It is zero when *p* = 0.5 and it is high when *p* = 0.25 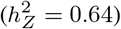 and tends towards 0 when *p* → 1 or *p* → 0 (not displayed on Figure 3). These results can readily be explained if we consider first that when one of the two alleles is rare, it is very unlikely that exactly two alleles of the same type interact to form a new collective at the next generation. For that reason, most collectives have a phenotype equal to 0, whether the parental phenotype is 0 or 1. Second when 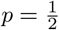, parental collectives, whatever their phenotype, always produce, on average, offspring collectives which all have the same collective-level character. This leads to 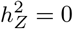. Finally, for values of *p* superior or inferior to 0.5 but not extremely superior or inferior (e.g., *p* = 0.25 and *p* = 0.9), collectives with *Z* = 1 tend to produce collectives which have a higher *Z* value on average than collectives with *Z* = 0. This is because under random assortment, and with these frequencies of alleles, it is more probable that parental collectives *AAaa* produce an offspring collective with the genotype *AAaa* than it is for any other parental genotypes. This results in a high collective-level heritability.

Although I do not show it here, the two conclusions that, firstly, in the absence of population structure (*f* = 0), collective-level heritability is always lower for non-linear traits that depend on the composition of collectives when compared to linear traits in the absence of population structure and, secondly, that it can be highly context-dependent (the more non-linear, the more context-dependent), can both be extended to larger collectives and different collective genotype-phenotype mappings.

## 5 Increasing Collective-level Heritability from Population Structure

Let us sum up what has been achieved so far. In Section 2, I showed that collective-level heritability of a trait that depends linearly on the trait of its constituent particles (which reproduce asexually and perfectly) is always equal to the heritability of the particle trait in the population of particles (which is equal to one), no matter what the population structure is. In Section 3, I argued that one important aspect of individuality is the ability for an individual to produce offspring that are on average not too dissimilar from itself 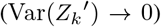. I then showed that one way to reduce the variance in offspring-collective character, given a parental-collective phenotype, is to increase the population structure (*f* > 0), so that the offspring particles produced by a parental collective have more chances to form an offspring collective together than with particles produced by other parental collectives, and consequently more chances to resemble their parental collective(s). In Section 4, I showed that when it comes to non-linear traits, even in very simple structures of two or four particles, the conclusion reached in Section 2, that collective-trait heritability is the same as particle-trait heritability, does not hold anymore. Collective-level heritability is always inferior to particle-level heritability and is highly context dependent, by which I mean that it varies widely with different frequencies of alleles in the population.

In this section I show that the conclusion reached in Section 2, namely that population structure does not affect collective-level heritability is not valid when the collective trait is a non-linear function of particle-level traits. I show that when traits are non-linear, population structure has two effects on collective-level heritability. First, insofar as population structure permits collectives to produce offspring with a lower collective-trait variance, it increases the heritability of non-linear collective traits. Second, it makes collective-level heritability less dependent on the general-population frequencies of alleles. Effectively, population structure has the effect of ‘linearizing’ non-linear collective traits by making the interaction of particles produced by a given collective less context dependent than when there is no population structure. In fact when *f* increases, because there is less shuffling between the particles produced by different collectives, the variability of the context of formation of offspring collectives decreases. This makes the particles of a collective effectively increasingly behaving as a single allele with a single effect.^9^

To see this, suppose we are dealing with our previous model in which a collective with four particles composed of two particles with allele *A* and two particles with allele *a* leads to a collective phenotype *Z* = 1, while all other genotypes lead to *Z* = 0. I do not present the two-particle case, but the conclusion reached with the four-particle case can be extended to collectives of any size. Suppose now that we have a population structure resulting in 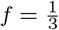. This means that offspring collectives have two parents, which both contribute two particles to each of their collective offspring. In this situation, like with the case presented in the previous section, to calculate collective-level heritability of character *Z*, we need first to know the average offspring-collective character of a given parental collective genotype. The collective character value (*Z*) for each possible collective genotype is reported in Table 6.

To calculate the average offspring-collective values reported in Table 6, I have used the following procedure which is explained for the parental-collective genotype *AAAa*, but it applies for all five possible collective genotypes. Collectives *AAAa* can transmit the combination of alleles *AA* in 50% of cases and the combination of alleles *Aa* in the other 50%. Since the alleles are at the (generalized) Hardy-Weinberg equilibrium, it is equivalent to choose the two other alleles for the collective at random. This means that the combination *AA* will form an *AA*-*AA* (*AAAA*) collective with probability *p*^2^, an *AA*-*Aa* (*AAAa*) collective with probability 2*pq*, and an *AA*-*aa* (*AAaa*) collective with probability *q*^2^. Similarly, the combination *Aa* will form a *Aa*-*AA* (*AAAa*) collective with probability *p*^2^, a *Aa*-*Aa* (*AAaa*) collective with probability 2*pq*, and a *Aa*-*aa* (*Aaaaa*) collective with probability *q*^2^. Note that in this model, if we assume that more than four offspring particles are transmitted from a parental collective, we consider first that two two-particle contributions are formed synchronically from the collective parental genotype and transmitted to an offspring collective. This operation is then repeated for the next four particles produced, and so forth. An alternative model would be that all offspring particles are produced at once, *and then* they interact randomly to form pairs and are transmitted to the offspring collectives. This latter model, which I do not explore here, produces different results from the one presented here.

**Table 6:**
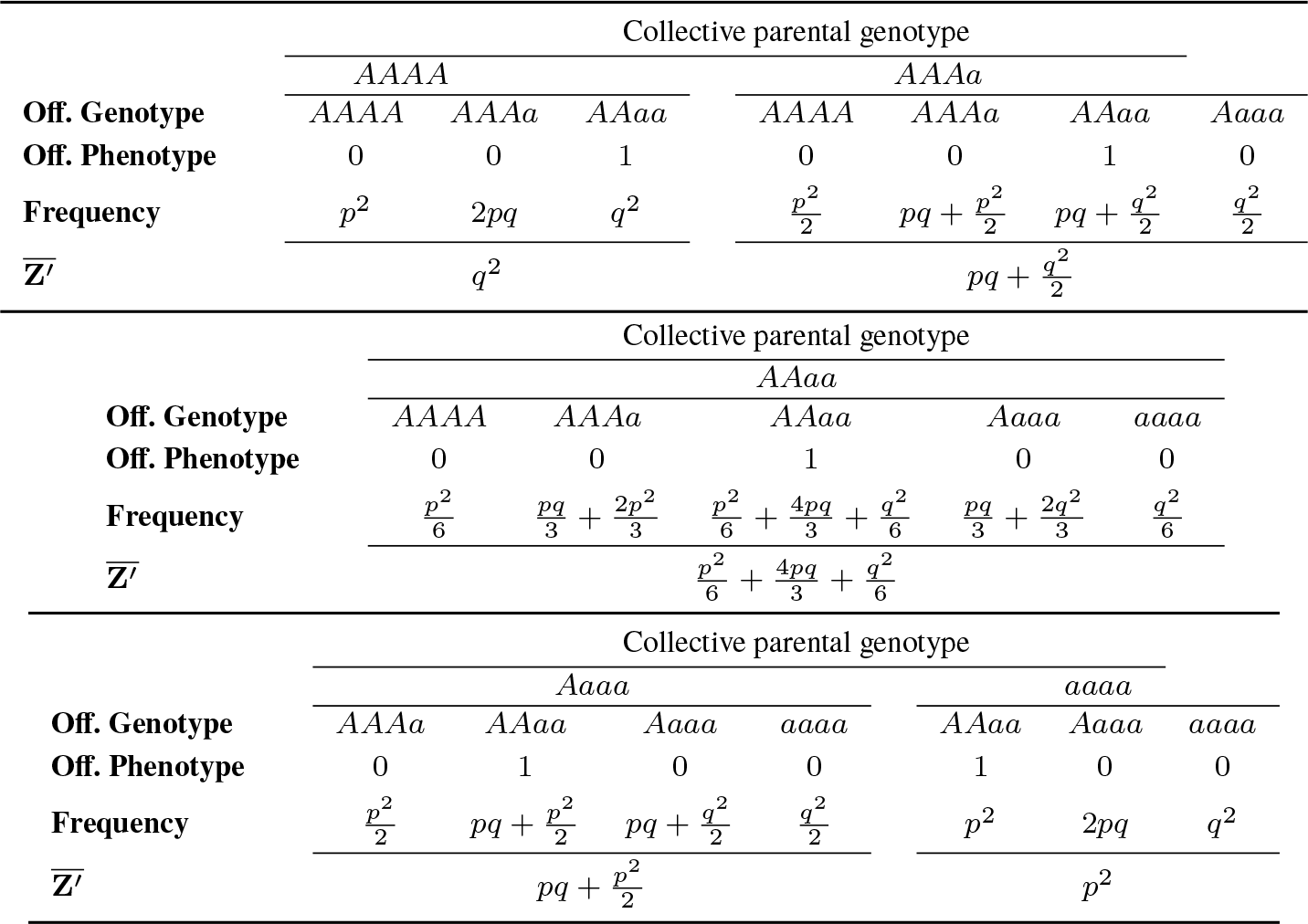
Frequencies of four-particle average offspring-collective phenotype for each parental-collective genotype with 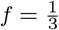

To have a direct point of comparison with the four-particle case when *f* = 0 discussed in Section 4, I assume that the four parental genotypes are at the generalized Hardy-Weinberg equilibrium, that is at the frequencies presented in Table 4. From Table 4 and Table 6, we can now calculate the weighted average offspring-collective character of parental collectives with *Z* = 0. It is given by the following equation:

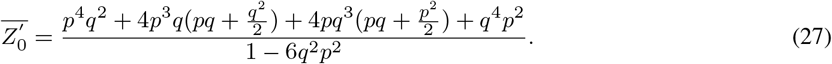

As with previous cases, 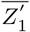 is found directly in Table 6 and is equal, in this particular case, to:

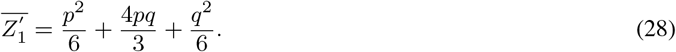

From there, like with the case where *f* = 0, we can now compute the linear regression of average offspring character on parental character, as shown in Figure 4. If we compare these results to the ones obtained in Figure 3, we can note that collective-level heritability varies less as the frequency *p* varies and is overall higher than when there is no population structure, even if for some particular values of *p* (e.g., *p* = 0.25) it is lower than when there is no population structure.

Let us now move on to the same four-particle case but with *f* = 1, that is a case where offspring collectives have only one parental collective. In such a case, as stated in Section 3, the variance in number of alleles transmitted from one parental collective to its offspring is nil whether the trait is linear or non-linear. In any case, the parental and offspring-collective characters are identical. The regression of average offspring character on parental-collective character is represented in Table 7.

In such a case, the frequency of parental collectives does not matter anymore to compute the parent-offspring regression. This is because the mean average offspring character is 0 for all collective genotypes with *Z* = 0 and 1 for all collective genotypes with *Z* = 1. This leads to the regressions of average offspring-collective character on parental-collective character shown in Figure 5 for the four frequencies of *p*, namely 0.25, 0.5, 0.9, and 0.99. When *f* = 1, collective-level heritability of a non-linear collective trait is, following my assumptions, one, that is equal to particle-level heritability. I have shown this result here with a four-particle collective case, but this can be extended to populations of collectives of any size.

**Fig. 4:**
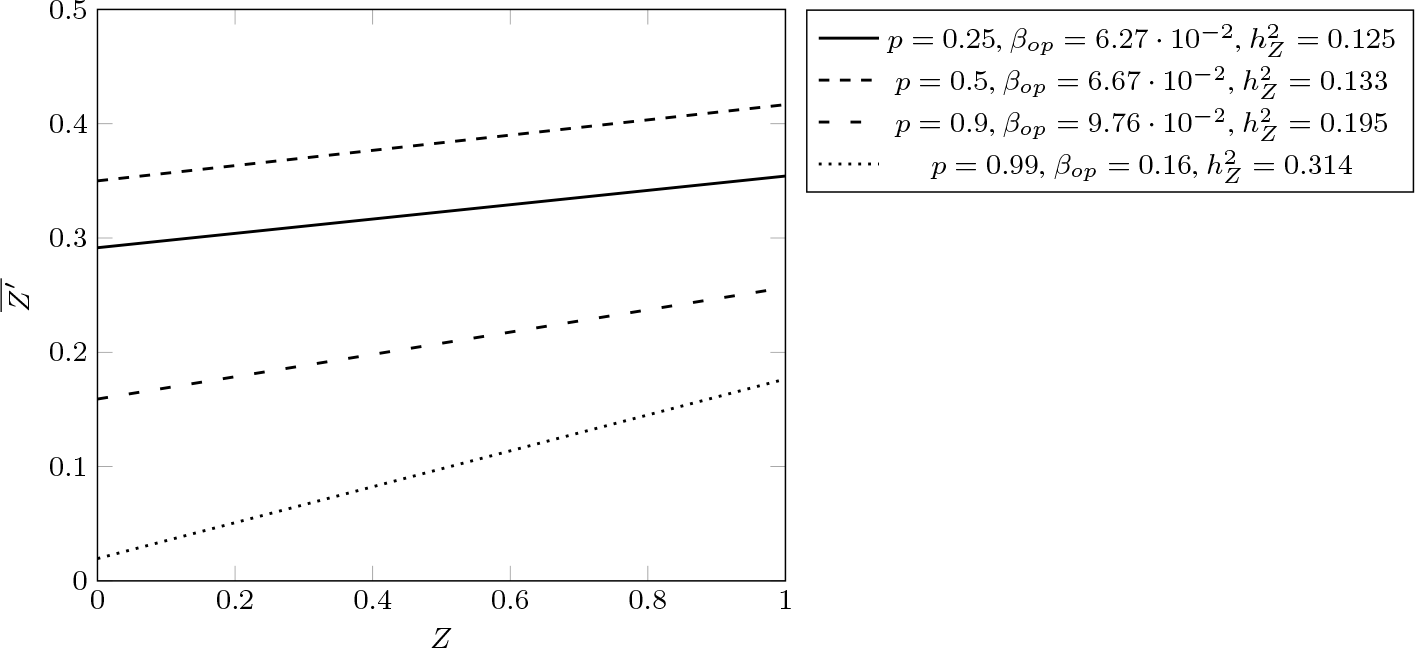
Linear regressions of average offspring-collective character on parental-collective character for the model of collectives with four particles producing a non-linear trait with moderate population structure 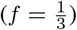. Collective-level heritability, in contrast to the cases in which there is no population (Figure 3) is less sensitive to the frequency *p* of allele *A*. *h*^2^ = 0.125 when *p* = 0.25, 0.133 when *p* = 0.5, 0.195 when *p* = 0.9, and 0.314 when *p* = 0.99.

**Table 7:**
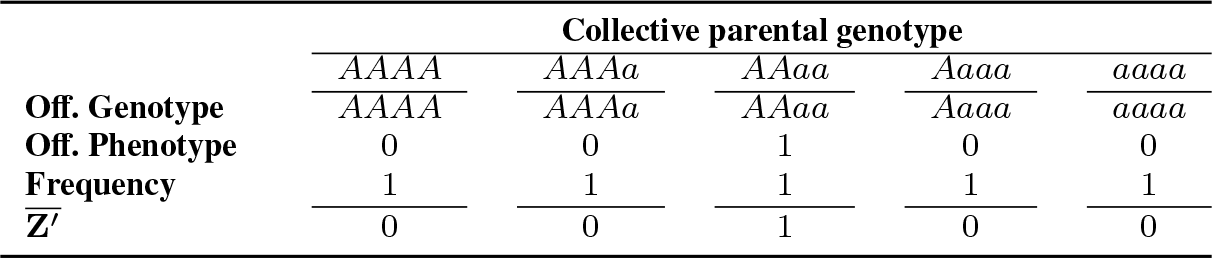
Frequencies of four-particle average offspring-collective phenotype for each parental-collective genotype with *f* = 1

**Fig. 5:**
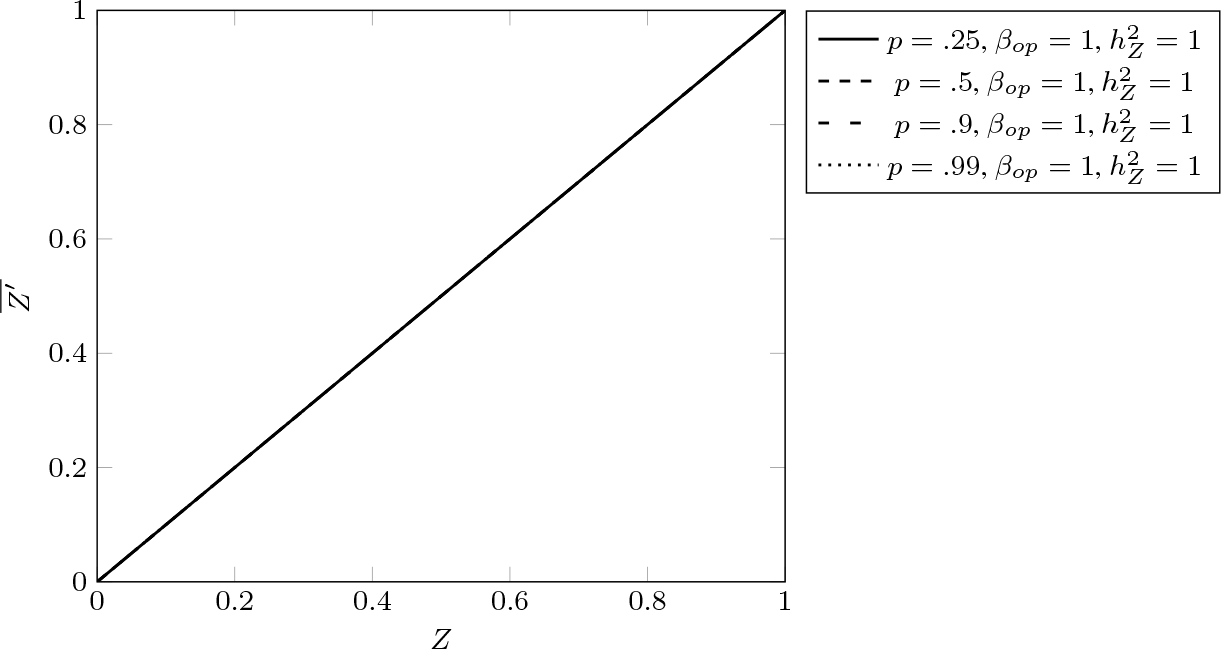
Linear regressions of average offspring-collective character on parental-collective character for the model of collectives with four particles producing a non-linear trait when population structure is maximal (*f* = 1). With maximal population structure, collective-level heritability is maximal (*h*^2^ = 1) and does not depend on the frequency *p* of allele *A*.

## 6 Discussion

With the different results derived in the previous sections in place, we can now ask their significance in the context of ETIs. I showed that when offspring particles interact randomly to form offspring collectives, collective traits that depend in a non-linear way on the proportion of particles within collectives can have a very small heritability for certain proportions of alleles in the general population. More importantly, variation in the frequency of alleles can change drastically the value of collective-level heritability. For instance, in the example with four-particle collectives, collective-level heritability varied from 0 when the frequency of *A* is 0.5 in the population, to heritability of around 0.36 when the frequency of *p* is 0.25 or 0.9, back to a very low frequency when the frequency of *p* is 0.99.

Such a huge variation in the value of collective-level heritability would make the long-term selection of a collective trait difficult. Imagine for instance a population of particles composed only of alleles *A* which interact randomly to form collectives. Suppose now that the new variant (*a*) emerges by mutation with a relatively low frequency. To take the four-particle case as an example, even if one collective was to exhibit two alleles *a* by chance and exhibit a collective phenotype *Z* = 1, that would confer a huge selective advantage, a low heritability at this frequency would mean that in spite of this huge advantage, its offspring collective would almost never tend to exhibit the same character.

What’s more, suppose now, for the sake of the argument, that a proportion high enough of allele *a* has arisen in the population, as a result of selection and/or drift, to a point where heritability is high (say *p* = 0.25). At that point, any change in the frequency of one of the two alleles, could easily lead to drastic change in heritability level. In a small population a change from *p* = 0.25 to *p* = 0.5 could readily happen. It would result in a collective-level heritability moving from almost 0.36 to 0. Such a huge variation would be an important obstacle for collective-level adaptation. In sum, when there is no population structure, in my example, having an advantageous collective-level character leads to a very unstable response to selection, which is unfavorable for collective-level adaptation to emerge.

If we now examine the effects of population structure, we can see that one of them is to increase collective-level heritability whether the frequency of *p* is high or low. For instance when *p* is 0.99, collective-level heritability moves from 0.06 when *f* = 0 to 0.314 when 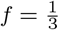. Furthermore, collective-level heritability is never lower than 0.12 when it could reach 0 when there is no population structure. This implies that the collective-level response to selection, when there is some population structure, is overall higher and less context dependent than when *f* = 0, that is whatever the frequency of the particle types (besides 0) in the global population is, there would always be *some* response to selection. From this, we can conclude that population structure acts as a buffer against variation in collective-level heritability. When *f* = 1, which is an extreme case, we can see that there is no context dependence of collective-level heritability: The four particles are always transmitted together. The non linear trait has effectively been ‘linearized’ so that the four alleles of each collective behave effectively as a single one. Another way to make this point is to say that non-additive genetic components of variance are ‘converted’ into additive genetic variance as *f* increases. I take the notion of ‘conversion of non-additive genetic variance’ from Goodnight (1988) (see also Wade, 2016, p. 12; Mackay, 2014) who has explored this phenomenon in the context of epistasis in small populations. I have shown here its importance for ETIs. When population structure exists or starts to increase, some traits that could not be reliably transmitted from parents to offspring at the collective level, so that no response to selection can occur at that level, starts to be reliably transmitted, and thus permits a response to selection to occur.

Although selection is not the main focus of this paper, it seems probable that population structure is a prerequisite for what one might call (advantageous) evolutionary innovations, that is phenotypes requiring the non-linear interaction between two or more particles for which there is no causal linear component of interaction, to be maintained over time. The collective non-linear trait presented in this and the previous section satisfies this definition. Population structure can in principle arise from different causal processes whether they are intrinsic or extrinsic to collectives. For instance, we could imagine two alleles at two different loci co-evolving, one conferring a direct evolutionary advantage by allowing the expression of a particular collective, while the second trait permits the linearization of the trait at that level. Another scenario could involve the existence of a pleitropic effect. As with two alleles, the two effects would be the same, but in this case one and the same allele would be responsible for both effects at once. A real case satisfying the two alleles scenario might be the evolution of extracellular matrix from cell walls permitting cells which were originally separating after mitosis to remain attached to one another (probably due to some genetic mutations). Herron (2017, p. 70-72) shows that different stages of cellular attachments exist in the volvocine algae lineage, with species in which extracellular matrix exists having arguably higher degrees of individuality than species in which no extracellular matrix is found. Given a mechanism of control of the number of cells per collective, the production of an extracellular matrix, by preventing cells to separate from one another, permits the reliable transmission of non-linear collective traits that would be impossible with freely moving cells. Finally, another causal origin could be ecology itself. In some suitable conditions, ecology could provide the template for stable collective realized heritability of collective-level traits to exist by preventing particles from different parents to interact, a hypothesis I explore elsewhere with collaborators (Black, Bourrat, Rainey, forthcoming).

The existence of population structure leading to a low variance in offspring character in the context of non-linear trait as a criterion for individuality is a novel one in the literature on biological individuality. Over the years many criteria have been proposed (for reviews see Clarke, 2010, 2013; Lidgard and Nyhart, 2017b; Pepper and Herron, 2008). Godfrey-Smith (2009, Chap. 5) (see also Godfrey-Smith (2013)), for instance, argues like others before him (e.g., Dawkins, 1982; Maynard Smith and Szathmary, 1995; Huxley, 1912), that one criterion for individuality is the existence of a bottleneck between collective generations. As recognized by Godfrey-Smith (2015) himself, the bottleneck criterion can only account for *fraternal* ETIs, that is, in transitions where the different partners forming collectives are closely related phylogenetically (Queller, 1997). In the case of *egalitarian* transitions, that is, transitions in which the different partners or particles of a collective have different phylogenetic origins, extreme bottlenecks (one single cell) cannot be achieved because there is no possibility for one partner to ‘represent’, that is to say reproduce on behalf of, the other(s). In contrast, for fraternal transitions reproducing on behalf of other partners this is readily achieved since all particles have the same genetic material. My criterion in terms of offspring variance and its evolution due to population structure best applies to egalitarian transitions. It is thus an alternative to the bottleneck criterion (which applies best to fraternal transitions). This criterion could also be used, in complement to other criteria to adjudicate the debate over whether holobionts (a macrobe plus its symbionts) and other multispecies entities such as biofilms represent genuine biological individuals (see Ereshefsky and Pedroso, 2013; Clarke, 2016b; Bourrat and Griffiths, 2018; Skillings, 2016).

I briefly mentioned that one reason to use a criterion of individuality in terms of low variance in offspring character for a given parental entity was inspired by Maynard-Smith and Szathmáry’s proposal that ETIs are associated with new ways to transmit information. The rational underlying this criterion is further propelled by noticing that it is often claimed that individuals are entities with the ability of ‘like begetting like.’ Yet, The notion of ‘like begetting like’ can be ambiguous. In fact, a high heritability might involve entities producing new entities that are very different from their parent (assuming there is only one parent) but, *on average* the offspring character has exactly the same value as the parental one. In such a case heritability (in the absence of environmental variation) would be unity. But the same expression might be understood in a different way, namely as the ability for a parent to produce reliably offspring that have a similar character value. In the former case, we would thus have a high heritability with an unreliable channel of transmission between parents and offspring, while in the latter case, not only would heritability be high but the channel of transmission would have a high fidelity. It is this second notion of ‘like begetting like’ that my criterion captures.

It should be clear that I am not claiming that this is the *sole* criterion that counts for individuality, for, like with any other measure or criterion for individuality, it would lead us to consider that some entities, for which the individuality status is regarded by many as equivalent, would score very differently. For instance, asexual entities, if the measure of variance in offspring character was taken to be the only important criterion, would score higher than sexual organisms. I am only claiming that a low variance in offspring character produced by an entity is one indicator for individuality. For a discussion on the tension that exists in evolution between evolutionary factors that increase genetic heterogeneity and those that increase homogeneity see Wright (1931, p. 142-147).^10^ Another way to make the same point is that in a population exhibiting genotypic variance, high fidelity between parent and offspring necessarily implies high heritability, but the converse is not true. The difference between these two notions of inheritance has often been overlooked and, I believe, has been a source of confusion in the literature. For an example of the type of confusion I am talking about see the debate between Maynard Smith (1987a,b) and Sober (1987), in which they use notions of inheritance without being clear whether they refer to the first or second sense I have distinguished.

## 7 Conclusion

In this paper, I have clarified in what sense collective-level heritability plays an important role in the levels of selection debate, and more particularly for ETIs. The outcome of my analysis is that collective-level heritability of non-linear traits can only be substantial and non-context dependent when there is a high level of population structure during the formation of collectives. The importance of non-linear interactions has long been noted in the multilevel selection literature (for a review see Wade, 2016). At the same time, most multilevel selection analyses focus on linear collective traits (e.g., Okasha, 2006). I have shown here that the implications of non-linear interaction for multilevel *inheritance* are equally important. The time is ripe to move the literature on the emergence of individuality to non-linear interactions. My model has remained highly idealized with several unrealistic assumptions such as an infinite population size and the same fitness for the two alleles. Given the difficulties to derive analytic results in the context of nonlinear systems, the next step in this project will be to build agent-based simulations and explore these two parameters jointly with the other parameters discussed in this article, as well as increase the number of loci and alleles responsible for the collective trait.

## Acknowledgments

I am thankful to Matthew Herron, Michael Bentley, and two anonymous reviewers who provided useful feedback on previous versions of this manuscript. I am also thankful to the Theory and Method in Biosciences group at the University of Sydney and more particularly Stefan Gawronski who proofread the final manuscript. This research was supported by a Macquarie University Research Fellowship and a Large Grant from the John Templeton Foundation (Grant ID 60811).

## Footnotes

1 For a recent update of the view developed in Maynard Smith and Szathmary (1995), see Szathmary (2015).

2 Note that because I am interested in the evolutionary origins of individuality, by ‘individual’ I will mean throughout ‘evolutionary’ or ‘Darwinian individual’. For other definitions of individuality and organismality see Pepper and Herron (2008); Gilbert et al (2012); Lidgard and Nyhart (2017a); Godfrey-Smith (2013). Note also that there is some tension with the view that a unit of selection can be equated with Darwinian individuality. In fact one might consider that individuality ‘emerges’ at one level when not one but a large number of traits at that level exhibit differences in fitness and heritability, while Lewontin’s conditions are trait specific. I will put these problems to the side here and consider the two as synonymous.

3 Note that the conditions require nevertheless to be slightly amended to fit the specificities of different levels of organization in the context of ETIs. For attempts to amend them see for instance De Monte and Rainey (2014); Bourrat (2014, 2015a); Griesemer (2000).

4 The notion of context dependence is notoriously ambiguous (Godfrey-Smith, 1992; Lloyd, 1988, p. 69; Sober and Wilson, 1994, p. 539). By ‘context dependence’ in this article I will mean ‘independent from the particle-type frequencies in the global population.’

5 For a similar approach to mine, in which the authors analyse the heritability of ‘heterozygoty’ in the context of diploid sexual species in which variation in the environment is not considered see Nietlisbach et al (2016).

6 Recall that I assume that all particles produce the same number of offspring at each generation in an infinitely large population, so that I keep both selection (i.e., difference in fitness associated with differences in phenotype) and drift out of the picture here.

7 For the difference between the binomial and hypergeometric distributions see Wroughton and Cole (2013).

8 For a model based on the ‘wrinkly spreader’ strain of *Pseudomonas fluorescens* (see Rainey and Rainey, 2003; Hammerschmidt et al, 2014), in which a fitness trade-off between collective viability and fecundity is considered, see Rainey and Kerr (2010). The sort of trade-off I have in mind here is slightly different as it concerns fidelity of transmission and fertility.

9 The notion of allele used here is similar to the one presented in Lu and Bourrat (2018), that is following an evolutionary conception of the gene, not a molecular one.

10 Thanks to Jonathan Hodge for pointing out this reference to me.

